# RNASeq analysis of *Aedes albopictus* mosquitoes during chikungunya virus infection

**DOI:** 10.1101/367235

**Authors:** Ravikiran Vedururu, Matthew J. Neave, Mary Tachedjian, Paul R. Gorry, Jean-Bernard Duchemin, Prasad N. Paradkar

**Affiliations:** CSIRO Health & Biosecurity, Australian Animal Health Laboratory, Geelong, Australia; School of Applied Sciences, RMIT University, Bundoora, Australia; School of Health and Biomedical Science, RMIT University, Bundoora, Australia

## Abstract

Chikungunya virus (CHIKV), preferentially transmitted by *Aedes* mosquitoes, is an emerging pathogen around the world and causes significant morbidity in patients. A single amino acid mutation in the envelope protein of CHIKV has led to shift in vector preference towards *Aedes albopictus*, an invasive mosquito. Previous studies have shown that after infection, mosquitoes mount an antiviral immune response. However, molecular interactions during the course of infection at different tissues and time-points remain largely uncharacterised. Here we performed whole transcriptome analysis on dissected midguts and head/thorax of CHIKV (Indian Ocean strain) infected *Aedes albopictus* to identify differentially expressed genes compared with uninfected controls. For this, RNA was extracted at two days post-infection (D2) from pooled midguts and eight days post-infection (D8) from heads and the anterior 1/3^rd^ of the thorax. We identified 25 and 96 differentially expressed genes from the D2 and D8 samples respectively (p-value <0.05). Custom *de novo* transcriptomes were assembled for the reads that did not align with the reference genome and an additional 225 and 4771 differentially expressed genes from D2 and D8, respectively, were identified. Twenty-two of the identified transcripts, possibly involved in immunity, were validated by qRT-PCR. Interestingly, we also detected changes in viral diversity, as shown by number of mutations in the viral genome, with increase in number of mutations in the midgut compared with mammalian host (Vero cell culture), followed by reduction in the number of mutations in head and thorax at D8, indicating a possible genomic bottleneck. Taken together, these results will help in understanding Aedes *Albopictus* interactions with CHIKV and can be utilised to reduce the impact of this viral infection.

**Author Summary:** Chikungunya virus has caused several outbreaks around the world in the last decade. Once a relatively unknown virus, it now causes seasonal infections in tropical and some temperate regions. This change in epidemiology is attributed to vector switch from *Aedes aegypti* to *Aedes albopictus*, an invasive pest leading to spread and causing infections in temperate regions. Although recent research has identified mosquito factors influencing infections, our understanding of interaction between chikungunya virus and its vector is limited. Using whole transcriptome sequencing of chikungunya infected mosquitoes, we identified differentially expressed genes in the midgut and head and thorax, over the course of mosquito infection. We also detected changes in the viral genome during mosquito infection and a possible genetic bottleneck event with reduction in viral variants at the head and thorax region of mosquito in the later stages of infection. These results will lead to improving our understanding of mosquito-virus interactions with *Aedes albopictus* as a vector and in turn lead to development of novel disease control strategies.

## Introduction

Arboviruses, such as dengue, chikungunya and Zika viruses, cause a significant burden on public health systems worldwide. Transmitted by mosquitoes, these viruses can cause high morbidity and mortality, with dengue alone causing more than 300 million infections per year[1]. First identified and described in 1955 in a report on an outbreak at the border of Tanzania and Mozambique in 1952, chikungunya virus (CHIKV) is an enveloped, positive sense RNA virus belonging to the Alphavirus genus in *Togaviridae* family[2, 3]. CHIKV infection causes high febrile illness, cutaneous exanthema and debilitating and often prolonged arthralgia [4–6].

While *Aedes aegypti* is the traditional vector for CHIKV, since the Reunion Island outbreak in 2005–2006, *Aedes albopictus* is observed to be more involved in viral transmission[7]. A single amino acid change in codon 226 of the E1 gene, that codes for the envelope protein of the virus, has improved fitness of CHIKV in *Aedes albopictus*[8]. This genetic shift has been implicated for most of the recent outbreaks, where despite an absence of the traditional vector *Aedes aegypti*, CHIKV has successfully established infections through *Aedes albopictus*. As an invasive species, *Aedes albopictus* has been expanding its traditional habitat of tropical and sub-tropical regions to much cooler temperate regions. *Aedes albopictus* also survives in favourable microhabitats even in winter and freezing temperatures [9]. These factors have further increased the risk of CHIKV to cause outbreaks in newer areas where mosquito-borne viral diseases are uncommon, such as Northern America and temperate Europe [10, 11].

For a mosquito to become infective, the virus needs to cross 2 critical barrier tissues, the midgut and salivary glands. Infection barriers can be influenced by multiple factors, including viral factors such as viral glycoproteins, or vector factors such as presence of a viral receptor, host replication factors and the microbiome composition of the midgut. Inside mosquitoes, after feeding, the blood meal moves down to the midgut where virus must contact epithelial cells before digestion of the blood meal and formation of the peritrophic matrix takes place. Following the successful infection of the midgut, the virus must overcome midgut escape barriers to disseminate to other tissues such as the haemocoel, head and salivary glands[12]. When an adult female *Aedes albopictus* mosquito is exposed to CHIKV in the process of blood feeding, the virus infects the midgut usually in a matter of hours [13–15]. From haemocoel, the virus makes its way to the salivary glands of the mosquito, which are present in the front 1/3^rd^ thorax region. Once the virus is detected in the saliva, the mosquito is considered to be infective and a competent vector [16, 17].

Although previous studies [18–23] have used transcriptome-based approaches to identify mosquito-virus interactions, tissue-specific responses during the course of infection have not been characterised. Here, using next-generation sequencing, we characterised the whole transcriptome response at the midgut (MG) (the first barrier site) and head & thorax (HT) (containing the salivary glands and viral dissemination sites), in *Aedes albopictus* in response to CHIKV infection. Our results also showed that the virus undergoes mutations in its genome as it passes through the mosquito and the number of mutations was significantly different at the two sites and in the course of infection, indicating a possible genomic bottleneck.

## Methods

### Chikungunya virus

Chikungunya virus isolate 06113879 (Mauritius strain), isolated from a viraemic traveller who returned to Australia in 2006, was obtained from the Victorian Infectious Diseases Reference Laboratory (VIDRL), Melbourne [24]. The isolate was passaged in Vero cells (Cell culture facility, AAHL) four times, followed by once in C6/36 (*Aedes albopictus* larval cell line, obtained from Cell culture facility, AAHL) followed again by Vero cells, which was then used for the experiments. TCID_50_ assay was performed on Vero cells to determine viral titer.

### Infections of *Aedes albopictus* and sample processing

All experiments were performed under biosafety level 3 (BSL-3) conditions in the insectary at the Australian Animal Health Laboratory, CSIRO. Insectary conditions were maintained at 27.5°C and 70% in relative humidity with a 12hr light and dark cycle. Female mosquitoes (5–8 days old) were challenged with a chicken blood meal spiked with CHIKV (1 in 100 dilution of stock virus, TCID_50_ 1.5 × 10^9^/ml) through chicken skin membrane feeding (Hemotek membrane feeding system®). Uninfected chicken blood and skin were provided by the Small Animal Facility (Australian Animal Health Laboratory) from chicken bred in the laboratory without any arboviral infection. The procedure was conducted with approval from AAHL Animal Ethics Committee. After one hour, the mosquitoes were anesthetised with CO_2_, blood fed females sorted and kept in 200 mL cardboard cup containers at 27.5°C, 70% humidity and 14:10 day:night photoperiod for 2 and 8 days with 10% sugar solution *ad libitum*. For controls, females were fed with blood mixed with media supernatant from uninfected Vero cell culture. Midguts were collected at 2 dpi (D2) and head/thorax were collected at 8 dpi (D8), from infected and control mosquitoes. Dissected tissues were stored in 50 μl of Qiagen RLTplus buffer with 5–10 silica beads (1 mm) at −80°C.

### RNA extraction and cDNA preparation

Bead beating was performed on MP Biomedicals FastPrep −24™ homogeniser, 3 cycles, speed: 6.5 m/s, 45 seconds each cycle. RNA was extracted using the RNeasy™ kit (Qiagen Australia) and cDNA was generated by using random hexamers and Superscript-III reverse transcriptase (Thermo Fisher Scientific Inc. Australia) following the manufacturer’s protocols.

### qRT-PCR for infection check and RNASeq data validation

cDNA generated from the RNA extracted from the midguts of D2 pools and the carcasses of D8 pools were tested for CHIKV viral RNA using an in-house designed qRT-PCR (Table B in S1), using primers specific for the E1 gene (Table A in S1). For RNASeq data validation, adult female mosquitoes were infected with CHIKV as described before and RNA was extracted from the MGs and HTs of 5 infected mosquitoes and cDNA was generated by protocols described previously. cDNA from the corresponding tissue of 5 uninfected mosquitoes was used as controls. qPCR was performed using gene-specific primers and 18s rRNA specific primers as internal controls (Table A in S1).

qPCR was performed on an Applied Biosystems Quantstudio™ 6 using the Takara-Clonetech SYBR Green Master Mix: SYBR Premix Ex Taq II (Tli RNase H Plus). The following cycling conditions were used with a melt curve at the end. Cycling conditions were: 30 seconds at 95°C, 40 cycles of 5 seconds at 95°C and 30 seconds at 60°C followed by a melt curve. The baseline and Ct values were calculated automatically using the supplied QuantStudio™ Software and the ΔΔCt values were calculated using the average ΔCt value of controls and 18srRNA as reference.

### RNASeq library construction and sequencing

Libraries for RNASeq were prepared using Nugen’s Ovation Universal RNASeq kit, following manufacturer’s specification with a minor modification in the HL-dsDNAse treatment. During first strand synthesis with DNase treatment, HL-dsDNase from Thermo Fisher Scientific was used in our library preparation, along with the 10x buffer supplied and their protocol used. The libraries were pooled and sequenced on a single lane of Hiseq-2500 (Macrogen Inc, South Korea) to generate 2 × 100 bp reads. The fastq files were deposited in NCBI’s Sequence Read Archive (SRA Accession ID: SRP140387).

### Data analysis-Differential gene expression and Gene ontology

Quality trimming of the raw sequences was performed using Trimmomatic v0.36. The reads were aligned to the CHIKV reference sequence (GenBank ID: MH229986) to assess the infection status using Hisat2 v2.0.5[25].

Following removal of *Gallus gallus* reads (due to chicken blood feeding) using SAMtools v1.3.1, the remaining reads were aligned to the *Aedes albopictus* Foshan strain genome sequence (AaloF1) from Vectorbase using Hisat2 and the resultant SAM file was sorted and converted into a BAM file using SAMtools[26, 27].

On Galaxy virtual lab v1.4.6.p5, featureCounts v1.4.6-p5 was used to quantify aligned transcripts from the sorted BAM files with default parameters for paired end reads, and DESeq2 v2.11.38 was used to obtain differentially expressed genes between the controls and infected samples by using default parameters on featureCounts output files [28, 29].

Using Trinity v2.3.2, two custom De Novo transcriptomes were built by combining the unaligned reads from MG (D2) and HT (D8) respectively[30, 31]. This transcriptome was used as a reference genome and the reads aligned, transcript counts measured and differentially expressed genes quantified using edgeR. The differentially expressed genes were annotated using BlastX and BlastN [32, 33].

The gene ontology (GO) IDs of the differentially expressed genes were obtained using the Biomart tool and topGO analysis was performed to identify the Molecular Functions (MF), Biological Processes (BP) and Cellular Components (CC) that were either enriched or depleted in the differentially expressed genes [34, 35]. Using topGO’s classic algorithm and based on p-values generated using Fisher’s exact method, differentially expressed genes were grouped based on their ontologies. Enrichment % was calculated as the ratio of the number of times particular genes in the pathway were differentially expressed compared to the expected number by chance.

### Whole genome sequencing of CHIKV

The Qiagen QIAseq FX Single Cell RNA Library kit was used for Illumina library preparation from total RNA extracted from CHIKV infected Vero cell culture supernatant using the Qiagen RNeasy™ kit. The library was sequenced on the Illumina Miniseq, with the mid-output kit (300 cycles) generating 2 × 150 bp paired-end reads. The resultant fastq files were quality trimmed and assembled to consensus sequence on CLC Genomics workbench v9.5.2. The sequence was annotated and submitted to GenBank (MH229986).

### Viral mutation calling and genetic diversity

From the infected D2 and D8 RNAseq libraries, the CHIKV reads were extracted using SAMtools. These reads were aligned to the reference sequence generated from the whole genome sequencing of the original cell culture isolate of the virus. Variant calling (SNPs and INDELs >3%) was performed using Varscan2 v2.3.9 on individual libraries as well as all D2 libraries merged together and all D8 libraries merged together [36, 37].

## Results

### RNASeq

For whole transcriptome analysis of *Aedes albopictus* mosquito tissues, during the course of CHIKV infection, dissected mosquito midguts at 2 dpi (as infection and first barrier site) and head/thorax at 8 dpi (as dissemination site containing salivary gland) were selected. The amount of RNA that could be extracted from a single *Aedes albopictus* midgut (MG) or head/thorax (HT) was below the lower limit of detection by the high-sensitivity kit on the Qubit (Thermo Fisher Scientific Inc. Australia). Hence, the MGs or HTs from 6 mosquitoes were pooled together for RNA extraction to obtain sufficient material. This was also required to avoid using a low-RNA input RNAseq kit, which would likely introduce bias during the PCR amplification stage. The pool size was kept as low as possible to retain information on biological variations.

To determine differentially expressed genes in MGs and HTs of *Aedes albopictus* mosquitoes infected with CHIKV, 5 pools of MGs collected at D2, and 5 pools of HTs collected at D8, from infected mosquitoes were created. For controls, 3 pools of MGs and 3 pools of HTs were used. Initially, to determine infection status of these tissues, qRT-PCR was performed using CHIKV-specific primers. Based on these results (Table C in S1) and requirements of the Nugen Ovation universal RNAseq kit, for D2, 2 control and 3 infected MG pools were used; while for D8, 1 control and 2 infected HT pools were used to prepare libraries.

The sequencing of eight libraries resulted in between 37 million and 170 million reads each. After quality trimming, reads mapping to the chicken genome were discarded to remove reads originating from undigested chicken blood. The remaining reads were aligned to the *Aedes albopictus* reference genome with the average alignment of 62.22% between the 8 libraries. The results also confirmed that all five infected libraries (3 from D2 and 2 from D8) contained viral reads, while the 3 control libraries (2 from D2 and 1 from D8) did not (Table 1).

**Table 1:** RNASeq NGS data summary. Total reads obtained, their alignment percentage to the reference genome and the percentage of reads aligned to the chikungunya genome.

|  | Total reads<br>(millions) | % mapped to<br>Refseq genome | % of CHIKV reads |
| --- | --- | --- | --- |
| D2_Infected MG 1 | 170.65 | 60.83% | 0.01 |
| D2_Infected MG 2 | 37.13 | 60.46% | 0.02 |
| D2_Infected MG 3 | 38.02 | 65.28% | 0.10 |
| D2_Control MG1 | 44.77 | 61.53% | 0.00 |
| D2_Control MG2 | 44.72 | 63.15% | 0.00 |
| D8_Infected HT1 | 47.15 | 63.28% | 1.12 |
| D8_Infected HT2 | 40.60 | 60.35% | 2.09 |
| D8_Control HT1 | 42.10 | 62.91% | 0.00 |

### Differential expression and TopGO analysis

Differentially expressed genes were identified using DESeq2 and edgeR and were plotted as Volcano plots (Fig 1). The results showed a number of genes to be differentially expressed at the two time points and tissue sites, using both methods (Table 2). The complete list of genes and transcripts differentially expressed with p-values of less than 0.05 is provided in supplementary information S2. The fasta files obtained as output for the custom transcriptome assembly are provided in supplementary information S3.

**Fig 1.**
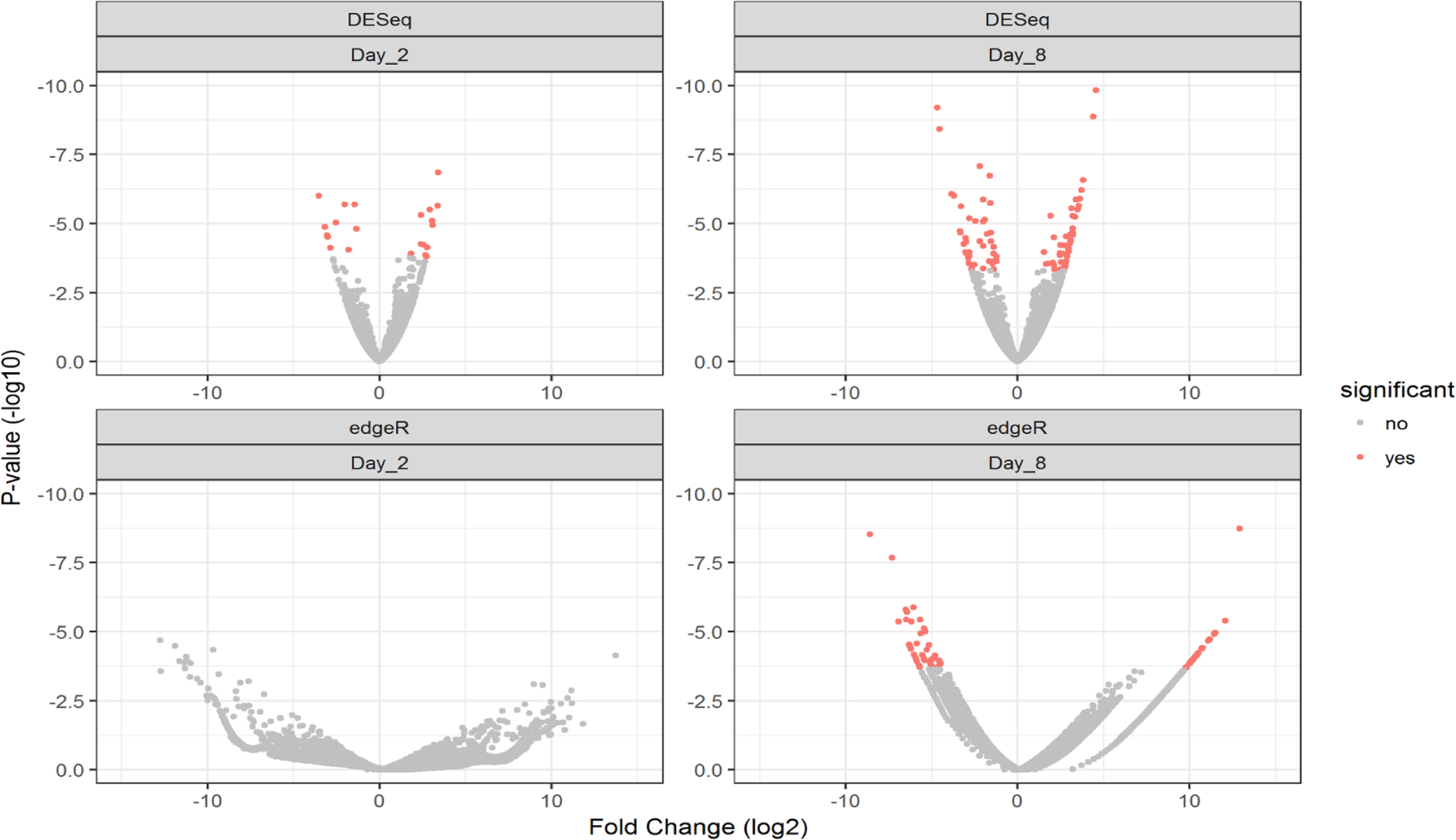
Volcano plots from DESeq2 and edgeR for D2 and D8 DGE. Volcano plots from DESeq2 (top panels) and edgeR (bottom panels) of differentially expressed genes from D2 (left panels) and D8 (right panels) samples. DESeq2 was performed by aligning reads to the *Aedes albopictus* reference genome; while edgeR analysis was done on reads that did not align to the reference genome and were aligned to the custom transcriptome.

**Table 2:** Differential Gene Expression analysis: Number of genes found to be differentially expressed in D2 midguts and D8 HTs using Deseq2 and edgeR analysis.

|  |  | No of Differentially expressed genes (P < 0.05) |  |  |
| --- | --- | --- | --- | --- |
|  |  | Total | Up | Down |
| D2 (MG) | DESeq2 | 25 | 14 | 11 |
|  | edgeR | 225 | 94 | 131 |
| D8 (HT) | DESeq2 | 96 | 51 | 45 |
|  | edgeR | 4771 | 3805 | 966 |

To determine the biological processes and molecular functions of the differentially expressed genes, gene set enrichment analysis and ontology was performed using TopGO (Fig 2). As expected, a number of biological and molecular processes were significantly affected during CHIKV infection.

**Fig 2.**
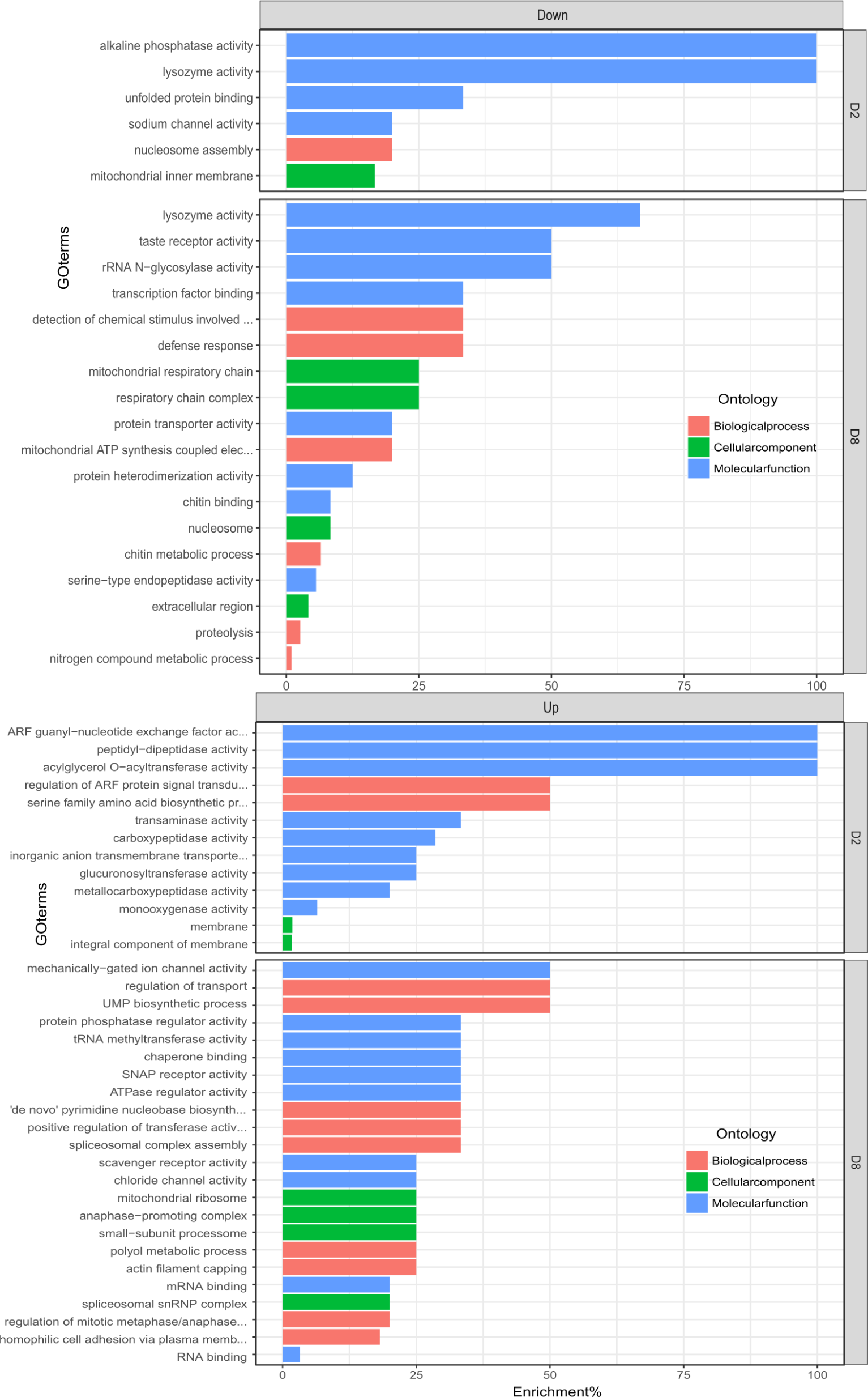
topGo enrichment comparison in differentially expressed genes. Enrichment analysis of down (left panel) and up (right panel) regulated genes in the midguts (D2) and heads and thorax (D8) of *Aedes albopictus* mosquitoes in response to CHIKV at 2 and 8 dpi respectively. Enrichment % is calculated as the ratio of ‘significant’ (Number of times the gene ontology number is observed as differentially expressed) to ‘expected’ (Number of times the gene ontology number is expected based on observation in control samples) gene numbers.

For D2 MGs, differentially regulated biological processes were metabolic (p-value 0.012) and serine family amino acid biosynthetic process (p-value 0.026). The differentially regulated molecular functions, likely of interest from an innate immune response point of view, were lysozyme activity (down regulated, p-value 0.028), alkaline phosphatase (down regulated, p-value 0.033) and carboxypeptidase activity (upregulated, p-value 0.042).

For D8 HTs, upregulated molecular functions were RNA (p-value 0.046) and mRNA binding (p-value 0.049) and molecular functions down regulated were lysozyme activity (p-value 0.0038), serine-type endopeptidase activity (p-value 0.0058), protein heterodimerisation (p-value 0.00181) and chitin binding (p-value 0.00532). Biological processes down regulated were defence response (p-value 9.70E-05), proteolysis (p-value 0.0075) and chitin metabolic process (p-value 0.0106). Up regulated biological processes were homophilic cell adhesion via plasma membrane (p-value 0.0056), UMP biosynthetic process (p-value 0.0245), regulation of transport (p-value 0.0245), spliceosomal complex assembly (p-value 0.0342).

Interestingly, genes involved in lysozymal activity were down regulated in both D2 MGs and D8 HTs, indicating significance of this pathway during infection as well as the dissemination process.

### RNASeq data validation on qRT-PCR

We selected 22 differentially expressed genes for validation by qRT-PCR, based on their known immune functions. This included 8 genes from D2 (1 aligned to reference genome and 7 from custom transcriptome) and 14 genes from D8 (11 aligned to reference genome and 3 from custom transcriptome) (see Table 3 for details).

**Table 3:** List of genes validated by qRT-PCR. List of genes selected for validation by qRT-PCR from the two time points and tissues and their annotation based on BlastX and BlastN.

|  | GeneID | Time point/<br>site | Gene Annotation |
| --- | --- | --- | --- |
| 1 | AALF020406 | D2/MG | Ankyrin repeat domain-containing protein 44 |
| 2 | TRINITY_DN109663_c0_g2_i1 | D2/MG | PREDICTED: mucin-22-like [*Aedes albopictus*]/<br>PREDICTED: flocculation protein FLO11-like, partial [*Aedes albopictus*] |
| 3 | TRINITY_DN110327_c1_g2_i1 | D2/MG | PREDICTED: ATP-dependent RNA helicase dbp2 [*Aedes albopictus*] |
| 4 | TRINITY_DN46100_c0_g1_i1 | D2/MG | PREDICTED: E3 ubiquitin-protein ligase MARCH6 [*Aedes albopictus*] |
| 5 | TRINITY_DN102975_c0_g1_i1 | D2/MG | PREDICTED: *Aedes albopictus* ARF GTPase-activating protein GIT2 |
| 6 | TRINITY_DN109582_c0_g2_i2 | D2/MG | PREDICTED: protein NPC2 homolog [*Aedes albopictus*] |
| 7 | TRINITY_DN110186_c0_g1_i2 | D2/MG | PREDICTED: translocon-associated protein subunit delta-like [*Aedes albopictus*] |
| 8 | TRINITY_DN110556_c0_g2_i1 | D2/MG | PREDICTED: uncharacterized protein LOC109408932 [*Aedes albopictus*] |
| 9 | AALF011899 | D8/HT | Quaking protein A |
| 10 | AALF004300 | D8/HT | PREDICTED: PDZ and LIM domain protein 7-like isoform X1 |
| 11 | AALF008354 | D8/HT | Putative ecdysteroid-regulated 16 kDa protein [Source:UniProtKB/TrEMBL;Acc:A0A023EHE1] |
| 12 | AALF012634 | D8/HT | PREDICTED: fat-like cadherin-related tumor suppressor homolog [*Aedes albopictus*] |
| 13 | AALF023547 | D8/HT | Peptidylprolyl isomerase [Source:UniProtKB/TrEMBL;Acc:A0A023EQ01] |
| 14 | AALF021910 | D8/HT | PREDICTED: fasciclin-2-like isoform X1 [*Aedes albopictus*] |
| 15 | AALF012324 | D8/HT | glycine rich RNA binding protein, putative |
| 16 | AALF016505 | D8/HT | leucine-rich immune protein (Long) |
| 17 | AALF025245 | D8/HT | PREDICTED: inhibitor of Bruton tyrosine kinase [*Aedes albopictus*] |
| 18 | AALF016704 | D8/HT | phosrestin i (arrestin b) (arrestin 2)<br>[Source:Projected from *Aedes aegypti* (AAEL003116) VB Community Annotation] |
| 19 | AALF026574 | D8/HT | Putative uncharacterised protein containing CCHC zinc finger domain |
| 20 | TRINITY_DN129476_c0_g1_i1 | D8/HT | PREDICTED: protein no-on-transient A isoform X2 [*Aedes albopictus*] |
| 21 | TRINITY_DN131737_c0_g1_i2 | D8/HT | PREDICTED: ficolin-3-like [*Aedes albopictus*] |
| 22 | TRINITY_DN131885_c0_g2_i2 | D8/HT | PREDICTED: PDZ and LIM domain protein Zasp-like isoform X5 [*Aedes albopictus*] |

The qRT-PCR results, using gene-specific primers, were compared with the RNAseq data (Table 4). In D2 samples, among the 8 targets chosen for validation, 6 were concordant with one target being discordant. In D8 samples, 7 out of 14 target genes were concordant. The ΔΔCt of TRINITY_DN46100_c0_g1_i1 (D2) and AALF008354 (D8), were less than 0.1 and hence not included in concordance calculation.

**Table 4:** Comparison of expression fold changes in chosen targets between qRT-PCR and RNASeq. Results of qRT-PCR on the 22 gene targets selected from RNAseq DGE analysis. Expression fold change for qRT-PCR is calculated as 2^-ΔΔCt and the up or down regulation is indicated by arrows, while LogFC was calculated by DESeq2 and edgeR

| Target ID | Expression fold change | LogFC |
| --- | --- | --- |
| AALF020406 | 0.5(↓) | -5.56 |
| TRINITY_DN102975_c0_g1_i1 | 0.76(↓) | -8.7 |
| TRINITY_DN109582_c0_g2_i2 | 5.35(↑) | 6.29 |
| TRINITY_DN109663_c0_g2_i1 | 0.16(↓) | -8.61 |
| TRINITY_DN110186_c0_g1_i2 | 1.88(↑) | 8.82 |
| TRINITY_DN110327_c1_g2_i1 | 6.53(↑) | -9.07 |
| TRINITY_DN110556_c0_g2_i1 | 1.51(↑) | 8.84 |
| TRINITY_DN46100_c0_g1_i1 | 1.03(↑) | -9.04 |
| AALF011899 | 0.8(↓) | 2.54 |
| AALF004300 | 0.43(↓) | 1.53 |
| AALF008354 | 0.96(↓) | 2.07 |
| AALF012634 | 0.01(↓) | 3.51 |
| AALF023547 | 0.75(↓) | 2.24 |
| TRINITY_DN129476_c0_g1_i1 | 0.01(↓) | -5.08 |
| TRINITY_DN131737_c0_g1_i2 | 0.07(↓) | -6.42 |
| TRINITY_DN131885_c0_g2_i2 | 2.9(↑) | 10 |
| AALF021910 | 2.97(↑) | 3.55 |
| AALF012324 | 0(↓) | 2.96 |
| AALF016505 | 3.65(↑) | -1.65 |
| AALF025245 | 17.54(↑) | 2.49 |
| AALF016704 | 1149.59(↑) | 1.51 |
| AALF026574 | 0.09(↓) | -3.12 |

The mean ΔCt values of D2 and D8 targets in infected mosquitoes compared to uninfected controls are shown in Table D in S1.

### Chikungunya viral variants in mosquitoes

Previous studies have shown that significant viral genetic bottlenecks occur during arbovirus infection of mosquito vectors [24, 38]. Each bottleneck event may result in significant reduction in viral variant diversity and thus affect the viral variant ultimately transmitted via mosquito saliva. To determine the CHIKV variant diversity in MG and HT samples, we first sequenced the CHIKV isolate 06113879. After passage in Vero cells, whole genome sequencing was performed using the Illumina MiniSeq, resulting in about 8.5 million quality trimmed paired-end reads. The assembly resulted in an 11,929bp long consensus sequence, which perfectly matched a previously sequenced 559bp portion of the E1 gene from this isolate (GenBank ID: EU404186.1).

The viral consensus sequence was most homologous (Identity: 11705/11985 (97.7%), Similarity: 11705/11985 (97.7%), Gaps: 245/11985 (2.0%)) to the CHIKV strain LR2006_OPY1 (GenBank: KT449801.1) and had the E1-A226V mutation, indicating this isolate also belongs to the same lineage as the CHIKV which caused the *La Reunion* outbreak in the Indian ocean in 2006.

Reads from the Vero isolate, as well as from D2 MG and D8 HT, were compared with the consensus sequence to determine the number of mutations in each sample. The number of mutations in the coding region of the D2 midgut samples (50; SNPs: 43 and Indels: 7) were significantly higher compared with Vero (16; SNPs: 9 and Indels: 7), while the number of mutations in D8 HTs were significantly lower compared with other samples (7; SNPs: 4 and Indels: 3). Interestingly, while most of the non-coding mutations were in the AT-rich and variable 3’ untranslated region (UTR), the Non-structural protein nsP3 gene was a hotspot for changes, particularly around position 5205 (Fig 3), indicating possible evolutionary pressures.

**Fig 3.**
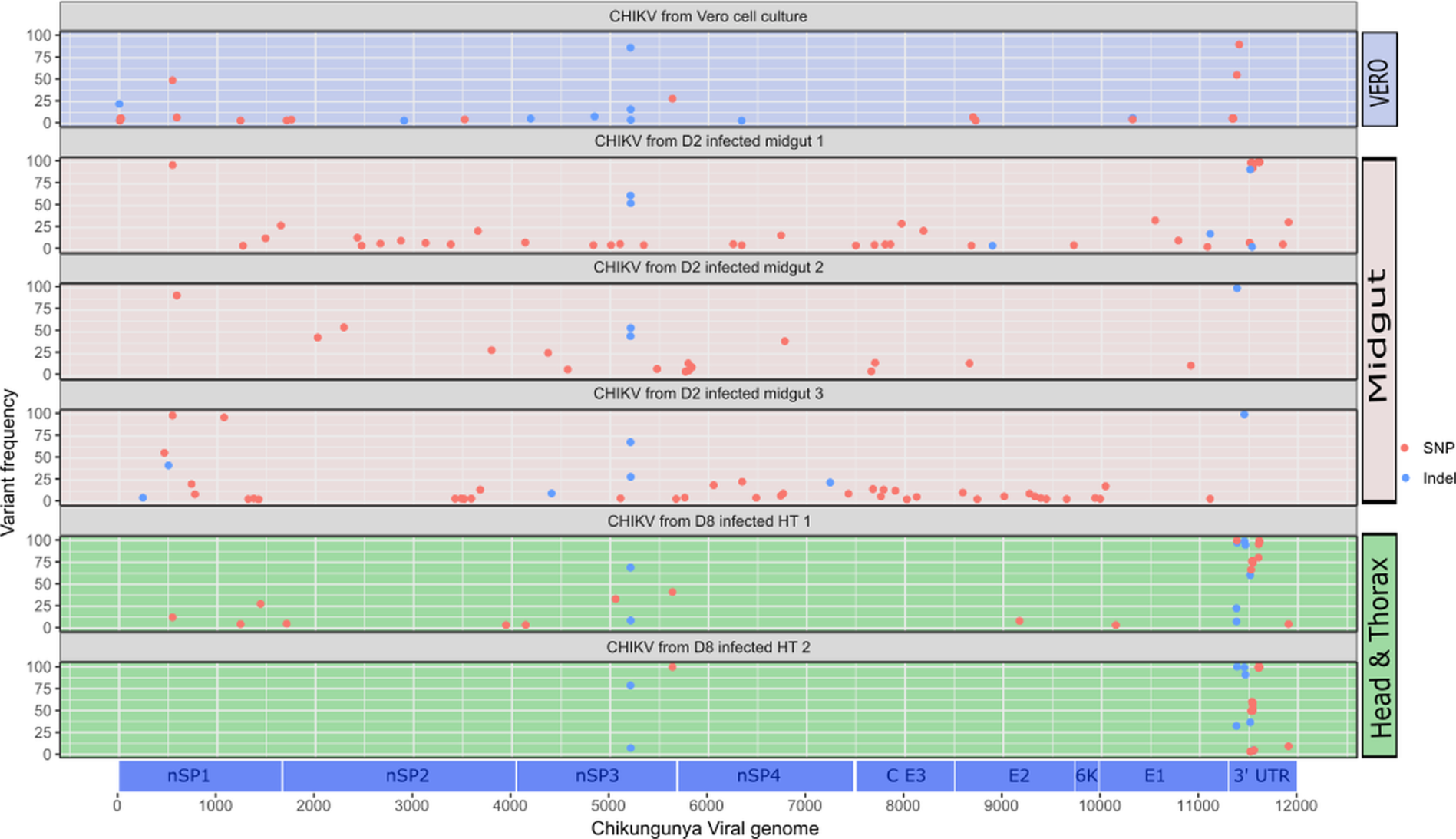
Variants detected in Vero cell culture isolate, D2 and D8 independent libraries plotted as per their location on the CHIKV genome. SNPs (red circles) and indels (blue circles) detected in the viral extract from Vero cell culture, three MG pools and two HT pools are plotted with reference to their position on the viral genome using VarScan2. The variants were called using Samtools and the figure was generated using R-Studio and annotated in Inkscape v0.91.

## Discussion

Chikungunya virus is a re-emerging alphavirus causing a high morbidity with long term arthralgia. Previous studies have taken approaches to understand the interaction between chikungunya virus and the *Aedes aegypti* vector [39]. However, considering the switch in vector preference towards *Aedes albopictus* by Indian Ocean isolates and invasive nature of this mosquito species, it is paramount to characterize the interaction between CHIKV and the new vector.

Previous studies with whole transcriptome analysis in mosquito vectors have used either whole mosquitoes or cell culture [18–22, 40]. Our objective here was to study the vector-virus interaction specifically at midgut, the first barrier site, and head/thorax, which denote the dissemination sites and include salivary glands, to understand the factors that play a critical role in determining mosquito vector competence.

In the current study, unbiased transcriptional analysis was performed on tissues collected from lab reared adult female *Aedes albopictus* mosquitoes over the course of CHIKV infection. The gene expression patterns at these tissue sites compared to uninfected samples revealed the transcriptional changes that are likely to be in response to the viral infection.

Our analysis revealed that at 2 dpi in midgut, most of the transcriptional changes are related to metabolism. Analysis of molecular functions revealed that while lysozyme activity and alkaline phosphatase were down regulated, carboxypeptidase activity was upregulated. Indeed, lysozymal and carboxypeptidase pathways are implicated in innate immune responses against multiple arboviral infections [41–44].

At 8dpi in head/thorax, differential regulation of biological processes including RNA and mRNA binding, lysosomal and serpin pathways and down regulation of defensin genes were observed. These could be due to either mosquito immune responses to CHIKV or viral modulation of immunity. Processes such as the regulation of transport and homophilic cell adhesion via plasma membrane could be involved in viral assembly and export [45–48]. Functional studies to determine whether these genes are pro or anti-viral are needed and could explain the role of these genes in the infection process.

Interestingly, in D8 HTs, two odorant binding proteins (OBPs) were found to be differentially expressed. Obp25 (AALF018602) was upregulated while D7 protein (AALF024478) was down regulated. In *Aedes aegypti*, salivary glands infected with dengue virus also showed differential expression of odorant binding proteins [49] and it was shown that down regulation of OBPs reduced the chemosensory abilities of the mosquitoes and hence reduced exposure to virus via feeding and thus hindered transmission capabilities. Similar mechanisms may be in play here as well, although that remains to be verified.

Multiple prior publications have also shown that the RNAi pathway is one of the major pathways involved in antiviral responses in insects [50–52]. In this study, we did not find any statistically significant changes in expression of genes involved in RNAi pathways. It is possible that the regulation of this pathway either does not occur at the transcriptional level or the proteins involved are ubiquitously expressed and not differentially regulated. It is also possible that the time points we selected did not coincide with RNAi activation.

Twenty-two genes were selected for validation by qRT-PCR, based on their possible involvement in mosquito immunity. The concordance was lower in the D8 genes compared to D2 genes. This could be due to the lower number of controls in D8 samples in the RNAseq analysis.

The incomplete and poorly annotated reference genome of *Aedes albopictus* was a hindrance in performing data analysis and robust pathway analysis. We also used heads and anterior 1/3^rd^ of the thorax, which included salivary glands at 8 dpi. The results from D8 samples represent data from heterogeneous tissue, and care needs to be taken before any broad conclusions are drawn. Functional characterisation of the identified genes may help in deciphering the results and understanding their role in mosquito-virus interactions.

RNA viruses, like CHIKV, have an inherent ability to rapidly mutate and generate variants for adaptation in novel environments through an error-prone polymerase[53]. Viral diversification is thought to be driven by the mosquito immune system leading to evolution of new genotypes. Previous study has shown that West Nile virus exhibits stochastic reductions in genetic diversity, which was recovered during intra-tissue population expansions [54]. Here, we showed that the mutations that arose in MG and HT samples compared to the original Vero cell culture isolate indicate possible changes in the viral sequence as it adapts from a mammalian host to different mosquito tissues. Vero cells, derived from African green monkey kidney, are known to be type-1 interferon response deficient [55], which may result in a high amount of CHIKV viral diversity as seen by high number of viral variants. Our results suggest that this diversity increases significantly in the mosquito midgut samples at 2dpi. We hypothesise that this increased genomic diversity represents CHIKV adapting and replicating in the mosquito midgut milieu. Interestingly, the number of CHIKV viral variants decreased considerably in HT by 8 dpi, possibly indicating a genetic bottleneck at this stage. These results are consistent with previous literature that show increase in CHIKV fitness and its adaptability during host switch [56]. The CHIKV 3’ UTR enhances viral replication in mosquitoes by interacting with mosquito cell-specific factors. This region contains highly conserved sequence elements for viral replication, binding sites for cellular miRNAs that determine cell tropism, host range, and pathogenesis, and conserved binding regions for a cellular protein that influences viral RNA stability [57]. The 3‘-UTR of the CHIKV genome showed the most variation in all our samples. Although significant, this may be due sequencing errors at this AT-rich region. Non-structural protein, nSP3, is an important protein for viral pathogenesis with immune modulatory functions [58]. We detected an apparent ‘hotspot’ of variability within the coding sequence of this protein in all samples, suggesting an important functional effect in mosquitoes that warrants further study.

Overall, our results showed significant changes in the transcriptome of *Aedes albopictus* mosquitoes after CHIKV infection, with identified genes involved in multiple cellular processes. This study, for the first time, examines differential gene expression at the midgut (the first critical barrier site) and head and thorax (dissemination site containing salivary glands) in infected mosquitoes. This study can be utilized in determining potential pro-viral and antiviral host factors and in turn, will be helpful in reducing the high impact of CHIKV infections by targeting the vector, *Aedes albopictus*.

## Acknowledgments

The chikungunya virus isolate was obtained from the Victorian Infectious disease reference laboratory, Melbourne. Thanks to Dr Kim Blasdell for helping with CLC Genomics workbench and to Chris Freebairn for collection of *Aedes albopictus* eggs from Torres Strait islands. Thanks to Adam Foord from AAHL-CSIRO for discussions about optimisation of qRT-PCR.

## Supporting information Captions

**S1 Supplementary information**. Table A: List of primers, Table B: Ct values of 10-fold dilutions of CHIKV RNA, Table C: Infection check of insect tissue pools and Table D: ΔΔCt of validated targets on qRT-PCR.

**S2 List of differentially expressed genes**.

**S3 Custom transcriptome output**.

